# STOmicsDB: a database of Spatial Transcriptomic data

**DOI:** 10.1101/2022.03.11.481421

**Authors:** Zhicheng Xu, Weiwen Wang, Tao Yang, Jing Chen, Yan Huang, Joshua Gould, Wensi Du, Fan Yang, Ling Li, Tingting Lai, Cong Hua, Shoujie Hu, Jia Cai, Honge Li, Lijin You, Wenjun Zeng, Bo Wang, Longqi Liu, Xun Xu, Fengzhen Chen, Xiaofeng Wei

## Abstract

Recent technological development in spatial transcriptomics allows researchers to measure gene expression of cells and their spatial locations at the almost single-cell level, which generates detailed biological insight into biological processes. However, specialized spatial transcriptomics databases are rare. Here, we present the Spatial TranscriptOmics DataBase (STOmicsDB), a user-friendly database with multifunctions including search of relevant publications and tools, public dataset visualization, customized specialized databases, new data archive, and online analysis. The current version of STOmicsDB consists of 141 curated spatial transcript datasets covering 12 species, and includes 5,618 spatial multi-omics publications and 674 tools. STOmicsDB is freely accessible at https://db.cngb.org/stomics/.

## Background

To understand cell development and biological functions, the gene expression profile of cells is a critical element (1, 2). Single-cell RNA sequencing (scRNA-seq) technologies characterize gene expression in single-cell resolution, which is a valuable tool for researchers to elucidate cell development. However, scRNA-seq loses spatial information of cells because tissues are dissociated during the experiment (3-5). By contrast, spatial transcriptomic technologies decode the gene expression of cells while retaining spatial information (5-7). This huge improvement allows researchers to analyze cell-cell interaction at the almost single-cell level. With the development of spatial transcriptomic technologies, especially the emery of high-throughput methods, such as 10x Genomics Visium (10x Visium) (8) and the recently released Stereo-seq (9), the studies based on spatial transcriptomic technologies are rapidly accumulating. Spatial transcriptomic technologies have been applied to many fields, such as disease research (10-14), organ atlases (4, 15-17), organogenesis (9, 18, 19), and plant biology (20-23). Due to the superiorities of the spatial transcriptomic technology in biological research, it was crowned Method of the Year by Nature Methods in 2020 (24).

With the increasing interest in spatial transcriptomics research, the lack of data archiving standards remains a big challenge. The major purpose of a data archiving system is to help other researchers to reuse and re-analyze the data. At present, most spatial transcriptomic data were deposited to Gene Expression Omnibus (GEO) in the National Center for Biotechnology Information (NCBI). However, GEO or other data repositories lack a spatial transcriptomic data archiving standard, resulting in various submission formats. A critical feature of spatial transcriptomics is the spatial information. For example, 10x Visium is the most common spatial transcriptomic technology. In general, 10x Visium has three types of information: gene expression data, the spatial information of barcodes, and histological images. Most submissions in GEO only included gene expression data, but lacked spatial information of barcodes or histological images. This absence of spatial information makes the reuse of spatial transcriptomic data challenging. Additionally, marker gene annotation and spatial variable gene annotation are useful for researchers, but this information is hardly provided in most GEO submissions. Another critical feature of spatial transcriptomics is that the data are obtained from tissue sections. A biological sample may be sliced into different tissue sections. Therefore, biological samples and tissue sections should be recorded during the data archiving. In sum, a specific data archiving standard for spatial transcriptomics is essential for researchers to reuse and reanalyze spatial transcriptomic data.

Towards the development of spatial transcriptomics, an online portal that integrates literature, tools, and data could provide invaluable support for researchers. However, these spatial transcriptomics databases are rare. SpatialDB (25) and Single Cell Portal (SCP) (26) are two spatial transcriptomics databases, which collect datasets in technology and study level, respectively. SpatialDB curated 24 spatial transcriptomic datasets across eight spatial transcriptomic technologies. It offers dataset visualization and comparison. However, the latest dataset that SpatialDB curated was in 2019, and SpatialDB has not collected datasets with the widely used 10x Visium technology. Furthermore, SpatialDB lacks a data archiving system. SCP, on the other hand, was initially designed for scRNA-seq data, but it has curated spatial transcriptomic data since last year. It archives data as studies. As SCP is a primary for scRNA-seq data, the spatial transcriptomic datasets are mixed with scRNA-seq data, which can only be retrieved by keyword search. As far as is concerned, around 20 spatial transcriptomic datasets were found by searching on 1 Feb 2022 on SCP. With the fact that the >150 public spatial transcriptomic datasets are available, and the amount is accelerated growing, SpatialDB and SCP only curated a small fraction of them. Furthermore, neither SpatialDB nor SCP offers interactional analysis between user data and database data, and none of them provide a specific spatial transcriptomic literature search and browse for users.

Here, we present Spatial TranscriptOmics DataBase (STOmicsDB), a user-friendly database serving as a one-stop service in the spatial transcriptomics field. STOmicsDB has five modules, the resource center module, the data exploration module, the customized database module, the online analysis module, and the data submission module. Firstly, the resource center module integrates 141 manually curated spatial transcriptomic datasets, and thousands of spatial multi-omics publications and tools for browsing and searching. Subsequently, the data exploration module provides comprehensive visualization and analysis of curated spatial transcriptomic datasets. Thereafter, the customized database module enables the collaboration with other researchers to construct specialized spatial transcriptomics databases. In the following, the online analysis module allows users to analyze their data with the datasets on STOmicsDB. Finally, the data submission module provides a spatial transcriptomic data archiving standard and archival system, allowing users to submit and deposit their data to STOmicsDB. In brief, STOmicsDB is the first spatial transcriptomics portal that provides analysis and visualization of existing datasets and comparative analysis of user data, a search of relevant publications and tools, and service of customized database construct and an archive of new data. It is anticipated that STOmicsDB could be served as an essential portal in the spatial transcriptomics field.

## Construction and content

### Data collection and curation

As described in the aforementioned context, one primary feature of STOmicsDB is the spatial resources collection and curation, which includes spatial transcriptomic datasets, and spatial multi-omics publications and tools.

To collect spatial multi-omics publications and tools, we firstly searched NCBI PubMed and PubMed Central with spatial multi-omics related terms to obtain candidates. Next, we manually curated hundreds of candidates to confirm whether they are related to spatial multi-omics. Subsequently, we used those curated candidates as the training set, and employed the machine learning method to further select and classify the rest spatial multi-omics publications. Subsequently, we manually checked each tool candidate to ensure that it is a spatial multi-omics tool. This resulted in 5,618 publications and 674 tools. We used automated scripts to retrieve the information of each publication or tool, which created the multidimensional and comprehensive metadata, such as included research areas, sample tissue, species, spatial resolutions, and publication types.

To collect the spatial transcriptomic dataset candidates, we retrieved the NCBI GEO and European Molecular Biology Laboratory-European Bioinformatics Institute (EMBL-EBI) ArrayExpress resources by searching the term ‘spatial’ and/or spatial transcriptomic technology terms, such as ‘MERFISH’ or ‘10x Visium’. Furthermore, we employed text mining on spatial multi-omics publications that curated before to search other spatial transcriptomic dataset candidates. Next, we manually curated each dataset candidate to confirm whether it is spatial transcriptome. We also included those spatial transcriptomics associated scRNA-seq datasets (under the same spatial transcriptomic project). This resulted in 85 spatial transcriptomic datasets and 9 associated scRNA-seq datasets. In addition, we collected other 45 spatial transcriptomic datasets from the 10x genomics website (8) and the SPATIAL research website (27). Finally, we curated 11 datasets which directly submitted to STOmicsDB. In the current version, they are deposited in China National GeneBank DataBase (CNGBdb) (28). In sum, the current version of STOmicsDB curated 141 spatial transcriptomic datasets.

### Data pre-processing and analysis

In brief, we used Scanpy (version 1.8.1) (29) to analyze curated datasets with default parameters. We firstly normalized and logarithmized the gene expression data from each dataset, and then we conducted principal component analysis (PCA) with the top 2000 highly variable genes to reduce the dimensionality of data. Next, we calculated the neighborhood map with PCA results. Uniform Manifold Approximation and Projection (UMAP) analysis and cluster spots were performed with the Leiden algorithm. For each dataset, we annotated cluster-specific marker genes with Wilcoxon rank-sum test by Scanpy. If a dataset contains spatial information, we then identified spatially variable genes using spatialDE (version 1.1.3) (30) with default parameters. Finally, we output the analysis results in h5ad format for data visualization.

### Data visualization

To visualize spatial transcriptomic datasets, we collaborated with the author of Cirrocumulus (31), and deployed this open-source application on STOmicsDB with customized modifications. Cirrocumulus was designed on rapid displaying large-scale scRNA-seq data and spatial transcriptomic data. It provides rapid and interactive data visualization.

### Database construction

We constructed the frontend framework with Vue.js (version 2.6.14) and built the backend using Django (version 2.2) and Python (version 3.7.4). STOmicsDB used PostgreSQL (version 9.6) to store the metadata of publications, tools, and datasets. We used Elasticsearch (version 7.16.2) as the search engine in the resource center of STOmicsDB. To manage and visualize curated datasets, we employed MongoDB (version 4.2) and Cirrocumulus. We used Redis (v5.0.4) as the cache to store and manage the data in memory. For task queue management, we applied RabbitMQ (v3.8.13). Nginx (v1.20.1) was used as the reverse proxy server. Currently, STOmicsDB supports following browsers: Google Chrome (v80.0 and up), Opera (v62.0 and up), Safari (v12.0 and up), and Firefox (v80.0 and up).

## Utility and discussion

### Overview of STOmicsDB

The current version of STOmicsDB curated 141 spatial transcriptomic datasets across 12 species, covering 19 spatial transcriptomic technologies. STOmicsDB also contains metadata of 5,500+ spatial multi-omics publications and 500+ spatial multi-omics tools. We provide general analyses and visualizations of most curated datasets.

STOmicsDB consists of five modules: the resource center module, the data exploration module, the customized database module, the online analysis module, and the data submission module (Figure 1). Users can access each module using the navigation bar on the top of the STOmicsDB home page (Figure 2).

**Figure 1.**
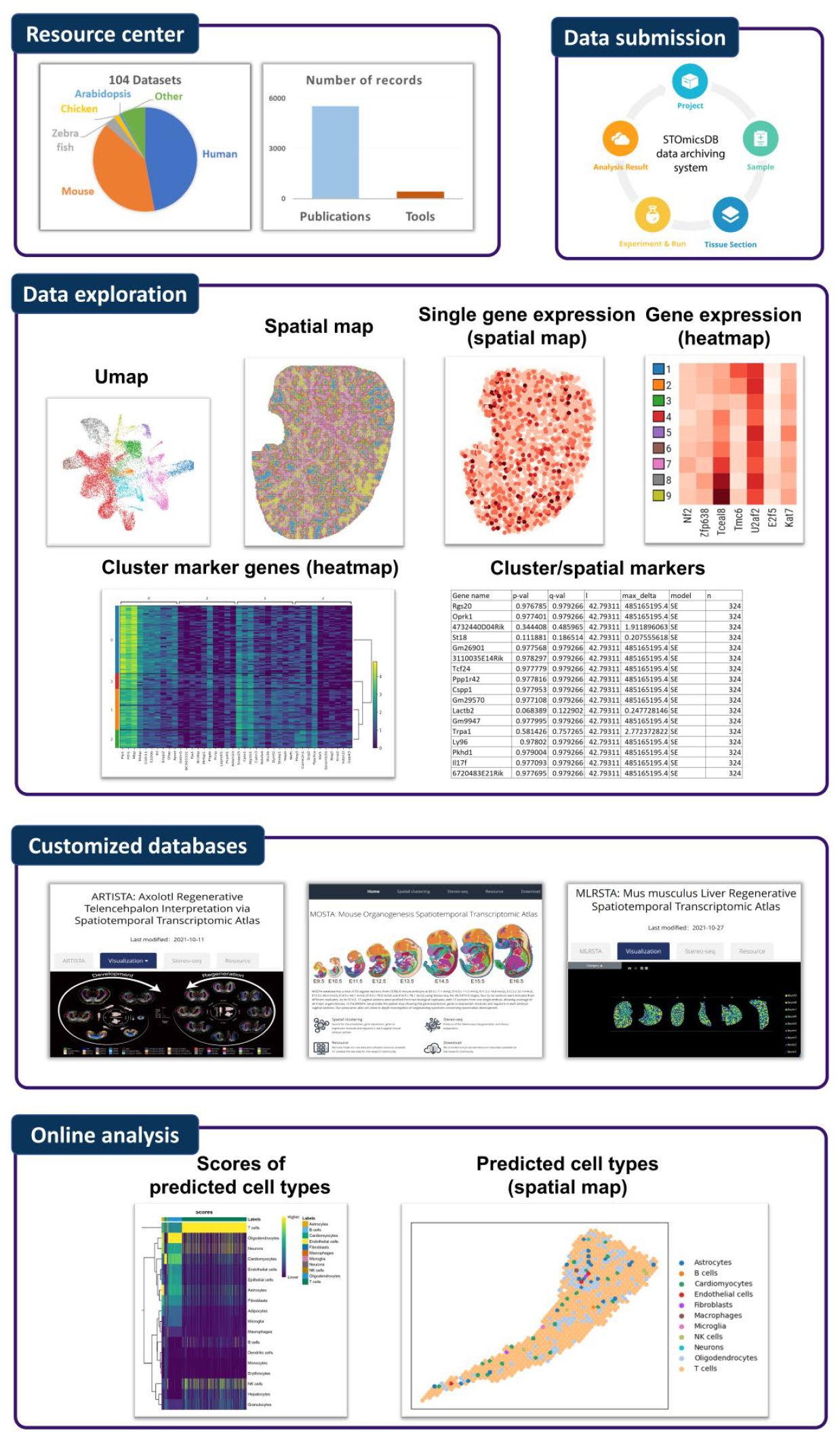
An overview of STOmicsDB.

**Figure 2.**
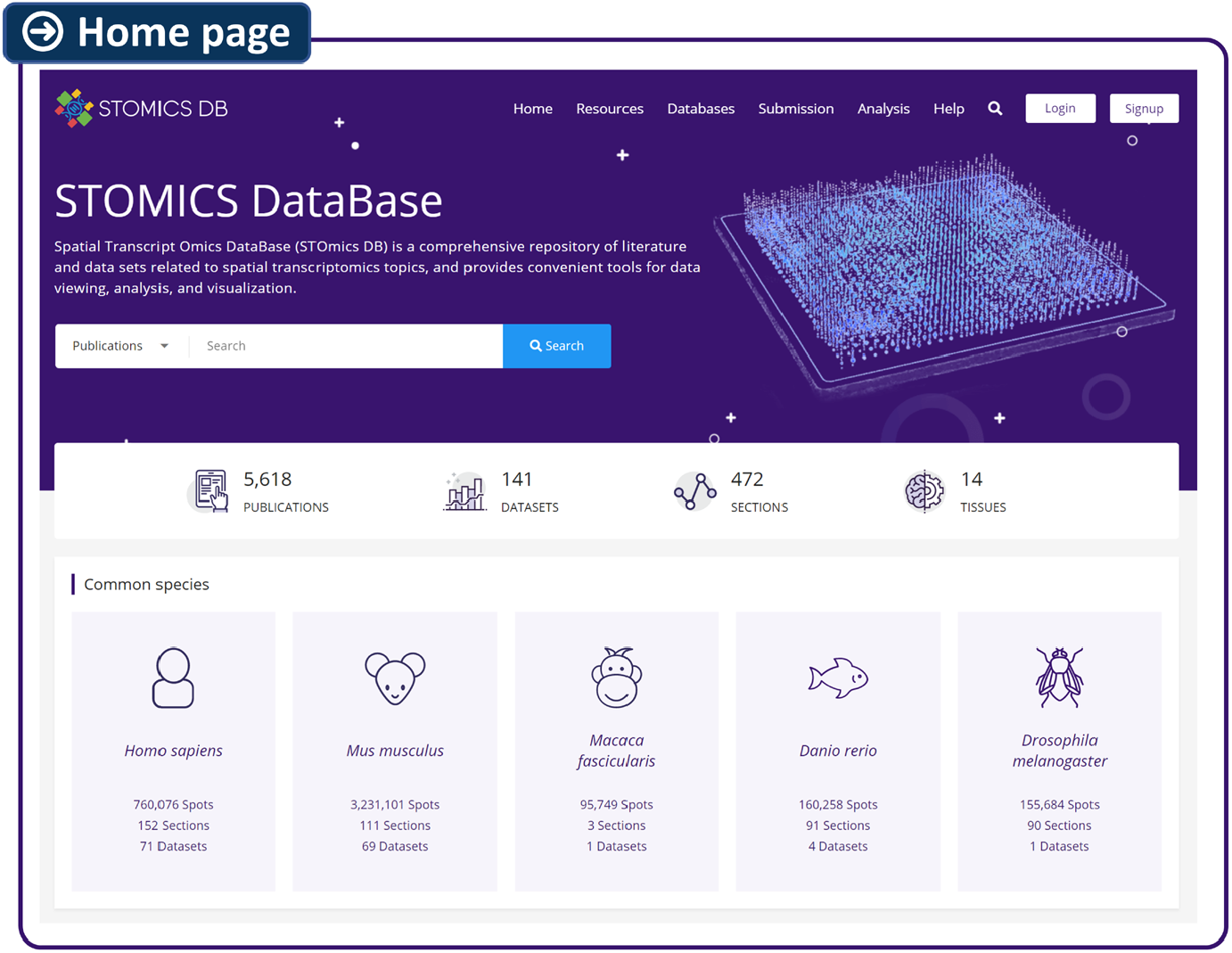
The interface of the home page of STOmicsDB.

### User-friendly resource center module

To meet the requirement for conveniently obtaining resources, STOmicsDB offers a comprehensive spatial resource center for searching and browsing, including three sections: (i) spatial transcriptomic datasets; (ii) spatial multi-omics publications; and (iii) spatial multi-omics tools. The spatial transcriptomic dataset section provides dataset visualization and analyzed data download, while the spatial multi-omics publication and tool sections contain the corresponding metadata and brief introduction of each record. Additionally, STOmicsDB have various classification for each section. For instance, the publication section contains research areas, species, tissues, spatial resolutions, and publication types (Supplementary Figure S1A), while the dataset section includes dataset release date, species, tissues, spatial transcriptomic technologies, and data quality (Supplementary Figure S1B). Users can specify the record of interest by ontology classifications. Furthermore, every record has an individual page, which displays detailed information, such as its summary and related links.

STOmicsDB provides two user-friendly searching methods in the resource center: a quick search and an advance search. For the quick search, a search box is presented on the home page, and users can select publications, tools, or datasets section using the drop-down list (Figure 2). For the advance search, it can be easily accessed by clicking the ‘Resources’ button in the top navigation bar. On the ‘Resources’ page, the left sidebar lists the section attributes (Supplementary Figure S1), which could be used as filter conditions. Users can download the metadata of interest by selecting the record and clicking the download button on the top right.

### Comprehensive data exploration module

As we continue curating public spatial transcriptomic datasets, STOmicsDB offers the opportunity for users to efficiently explore most of latest spatial transcriptomic datasets. To the best of our knowledge, STOmicsDB is the database that provided the most spatial transcriptomic dataset visualization (>100), compared with other databases, such as SpatialDB or SCP.

STOmicsDB provides two types of data exploration: (i) interactive data visualization; (ii) data analysis results. Users can visualize our curated datasets in the dataset section of the resource center, by entering the dataset page and selecting the ‘Explore’ tab or ‘Analysis results’ tab on the top left of that dataset page (Figure 3). Different section/sample in the same dataset can be chosen through ‘Sections’ selector on the top. For the interactive data visualization (‘Explore’ tab), STOmicsDB provides UMAP and spatial map (if the dataset contains spatial information) for each dataset, which can be switch using the ‘Embeddings’ on the sidebar. Users can define the color, name of each cluster, and move or zoom the interactive image. If users select a gene/genes, a gene expression heatmap (on UMAP or spatial map) will be displayed on the right, and a heatmap or violin plot of gene expression will be shown by selecting the ‘DISTRIBUTIONS’ button in the top toolbar. STOmicsDB also supports the expression comparison with two selected regions. In addition, STOmicsDB shows plots of data statistics, and figures and tables of cluster marker genes and spatially variable genes in the ‘Analysis results’ tab. This information helps users to efficiently obtain the corresponding marker genes. All plots and marker information can be downloaded.

**Figure 3.**
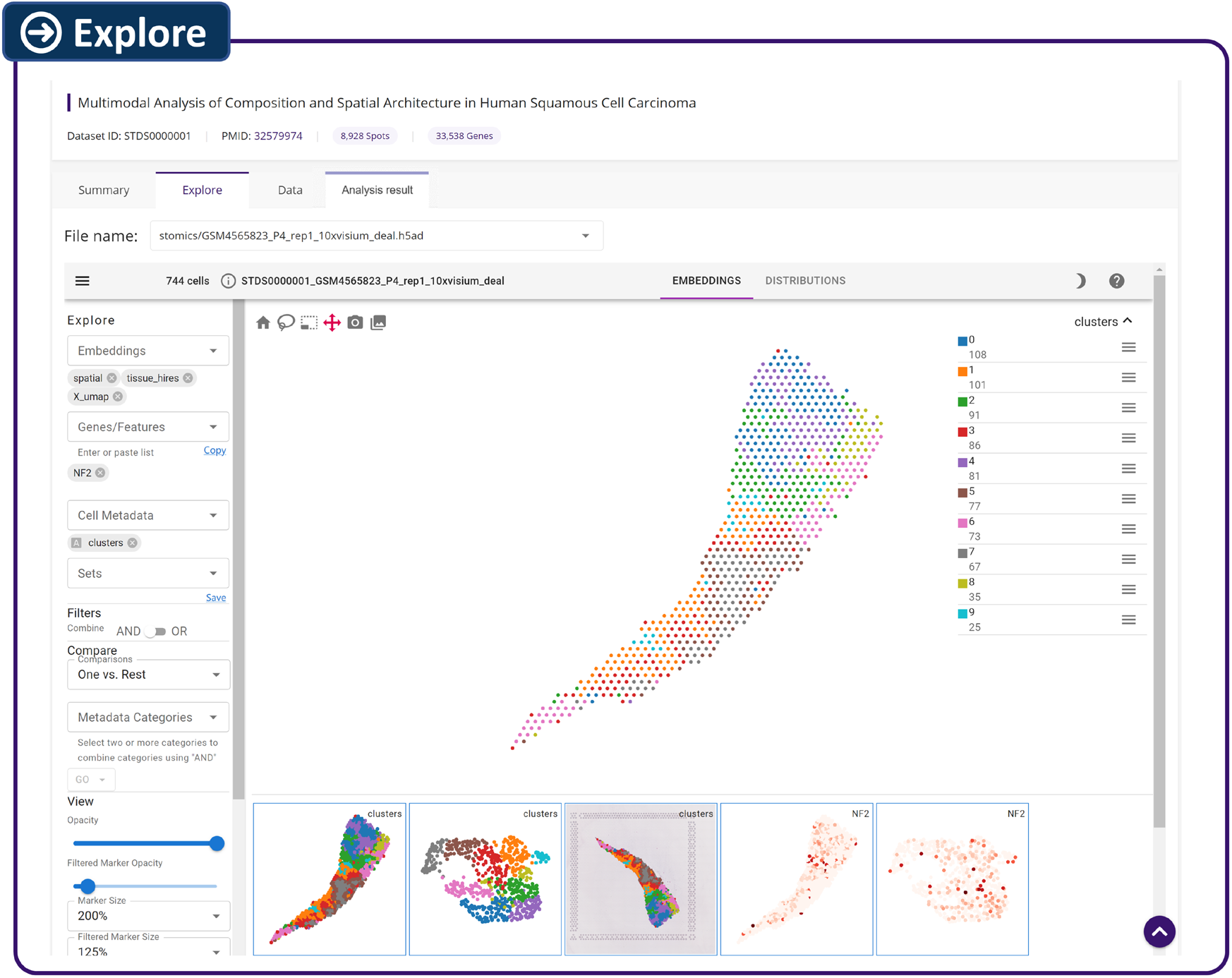
The interface of the visualization tool. The left sidebar is the parameter options, and the right part is the interactive image. The bottom left part displays history figures.

Here, we provided a user case to illustrate the potential usage and application of our data exploration module. For instance, the dataset STDS0000113 is related to kidney injure in human and mouse (32). We found that *Mettl7a2* is a marker gene of cluster 0 in GSM5224981 (kidney of cecal ligation puncture murine model) section in the cluster marker table. UniProt (33) suggests that *Mettl7a2* is related to methyltransferase activity, and methyltransferases may be associated with kidney injure (34, 35). We then checked the expression level of *Mettl7a2* by clicking “Mettl7a2” in the cluster marker table. The spatial map showed that *Mettl7a2* has spatial expression heterogeneity, which only has high expression in a specific region (Figure 4). As accurate cell type annotation is still challenging, we have not annotated each cluster to avoid introducing possible errors. The original research suggests that the specific region that *Mettl7a2* highly expressed could be proximal tubule (32). Proximal tubular cells are susceptible to be damaged during kidney injure (36, 37), and DNA methylation might protect the proximal tubular cells (38). It implies that *Mettl7a2* may pay a role in kidney injure in proximal tubule. In short, STOmicsDB provides opportunities for users to efficiently explore gene of interest in most published spatial transcriptomic datasets.

**Figure 4.**
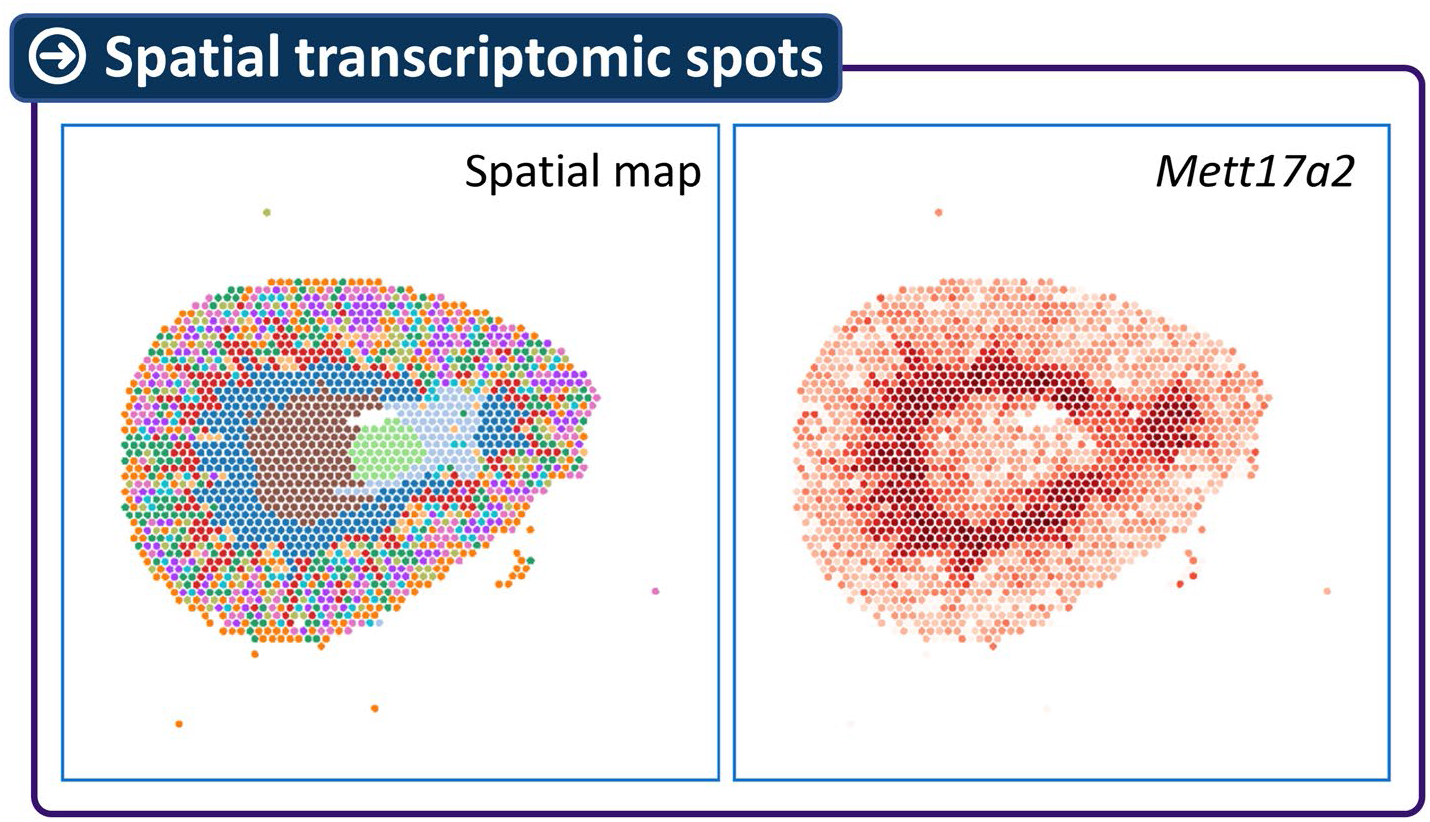
Spatial transcriptomic spots of kidney of cecal ligation puncture murine (a case study). Left: the spatial map coloring by unbiased clustering. Cluster 0 was colored by blue. Right: the expression of *Mettl7a2*.

### Customized database module

One significant feature of STOmicsDB is that STOmicsDB provides a customized database service. We collaborate with other researchers to construct specialized databases according to their spatial transcriptomic data and research purposes. Under this collaboration, researchers provide the data, and we work with them to set up the database structure and data visualization. Now, we have constructed three such databases with other researchers: ATRISTA (related to axolotl brain regeneration), MOSTA (related to mouse organogenesis), and MLRSTA (related to mouse liver regeneration) (Figure 1). Moreover, we also welcome database hosting. Researchers can construct the spatial transcriptomics database by themselves, and deploy the database on STOmicsDB. Users can browse these customized databases by clicking the ‘Databases’ button on the top navigation bar (Figure 2).

### Online analysis module

Compared with scRNA-seq, spatial transcriptomic experiments are expensive. To facilitate the use of spatial transcriptomic data, we set up an online tool based on SingleR (v1.4.1) (39) to provide an interactive analysis between user scRNA-seq data and spatial transcriptomic data that STOmicsDB curated. This tool allows users to annotate cell types of a specific spatial transcriptomic dataset on STOmicsDB by uploading their scRNA-seq gene expression matrix and the corresponding cell type annotation. Except for the default outputs of SingleR, this tool also generates a spatial feature plot to show the spatial localization of each annotated cell type (Figure 1). This is useful for users to obtain the corresponding spatial information about their scRNA-seq data with the help of the large curated spatial transcriptomic datasets on STOmicsDB.

### Data submission module

The lack of spatial transcriptomic data archiving standards makes data reuse and re-analysis challenging. We have developed the spatial transcriptomic data submission standard to overcome this obstacle. This spatial transcriptomic data submission system includes five parts: Project, Sample, Tissue section, Experiment & Run, and Analysis result (Figure 1). The Project, Sample, and Experiment & Run parts record project information, biological sample information, and related experiment information, respectively. These three parts are the same as traditional data archiving systems, such as Sequence Read Archive (SRA). Due to the feature of spatial transcriptomics, the data are generated from the tissue section, and each sample could have multiple tissue slices. We, therefore, included the tissue section information into our data archiving system. The final part of our submission system is the Analysis result part. Different technologies have different default analysis outputs. For instance, the positions of spatial spots are deposited into a text file by the output of 10x Visium technology, while Stereo-seq stores this information into a binary file. We developed different standards for different technologies according to their features to handle these different features (Figure 5). Additionally, STOmicsDB allows users to submit downstream analysis files, such as marker identification results, differential expression results, or cluster annotation results. In sum, our spatial transcriptomic data submission system archives almost all spatial transcriptomic data analysis results, from raw data to downstream analysis, which facilitates the spatial transcriptomic data reuse and further research.

**Figure 5.**
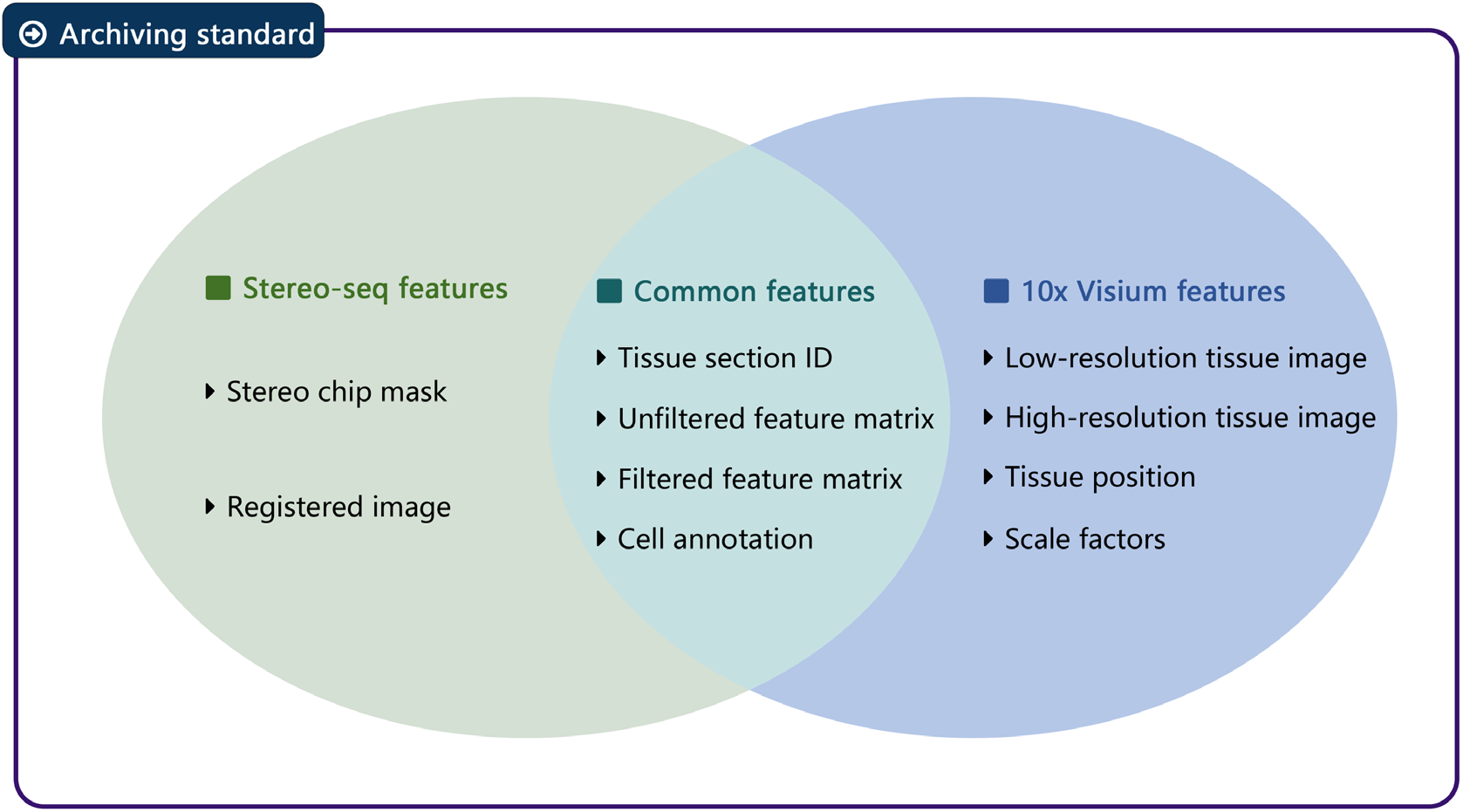
An example of technology-specific data submission standards. Each spatial transcriptomic technology may have specific features. STOmicsDB developed technology-specific standards to fit different conditions.

### Further developments

In the future, we will improve STOmicsDB in the following directions. Firstly, we continue curating datasets/publications/tools when numerous new spatial transcriptomics projects are being studied. Next, we plan to create multi-level interactions among publications, authors, tools, technologies and datasets. For example, these multi-level interactions will show which tool that most papers used, or how many spatial multi-omics relevant papers that a specific author published. This enables users easily obtain latest and comprehensive information on spatial transcriptomics field. Furthermore, a comprehensive online spatial transcriptomics submission system is invaluable. In the current version, the data archiving system on STOmicsDB only supports submission through command line, lacking a user-friendly visual interface. We are developing this interface, and will deploy it on STOmicsDB in the next version. Moreover, although we have collected and curated 19 spatial transcriptomic technologies, we still struggle to support an interactive visualization for every technology. Finally, we plan to integrate the marker genes of curated datasets of the same species or the same organ to generate gene networks or relevant atlases. These networks may improve our ability to efficiently characterize biological insight into cells and tissues.

## CONCLUSIONS

To our knowledge, STOmicsDB is the first portal that integrates multiple aspects of spatial transcriptomic resources, analyses, and services, which provides invaluable support for researchers in the spatial transcriptomics field. STOmicsDB has multiple novel features. Firstly, the research center in STOmicsDB curated hundreds of datasets and thousands of publications and tools, as well as comprehensive classifications of each record, allowing researchers to locate the resource of interest efficiently. Thereafter, the data exploration module in STOmicsDB offers researchers the opportunity to access, visualize, and analyze numerous spatial transcriptomic data, saving researchers’ time for public data collection and analyses. Furthermore, the customized database module in STOmicsDB supports researchers in constructing specialized databases, assisting in comprehensive data visualization. In the following, the online analysis module in STOmicsDB allows researchers to explore their data with a wide variety of curated spatial transcriptomic data on STOmicsDB. Finally, the data submission module in STOmicsDB first developed a specific spatial transcriptomic data archiving standard, which addressed the data reuse issue. In sum, it is believed that STOmicsDB serves as a one-stop service in the spatial transcriptomics field, which would be of great benefit to the spatial transcriptomic community.

## Abbreviations

CNGBdb: China National GeneBank DataBase
EBI: European Bioinformatics Institute
EMBL: European Molecular Biology Laboratory
GEO: Gene Expression Omnibus
NCBI: National Center for Biotechnology Information
SCP: Single Cell Portal
scRNA-seq: Single-Cell RNA sequencing
STOmicsDB: Spatial TranscriptOmics DataBase
SRA: Sequence Read Archive

## Declarations

### Ethics approval and consent to participate

Not applicable.

### Consent for publication

Not applicable.

### Availability of data and materials

The analysis data generated during the current study are available at https://ftp.cngb.org/pub/SciRAID/stomics/.

The datasets used during the current study are available at CNGBdb (https://db.cngb.org/) with accession number: CNP0001543, CNP0001469, CNP0000927, CNP0002189, CNP0002220, CNP0002068, CNP0002199, CNP0002310, CNP0001923, CNP0002316, CNP0002347, CNP0002110, CNP0002590, CNP0002543, CNP0002646, CNP0002476, CNP0002618; NCBI GEO (https://www.ncbi.nlm.nih.gov/geo/) with accession number: GSE65924, GSE104292, GSE111672, GSE114723, GSE114770, GSE114770, GSE116008, GSE118403, GSE120374, GSE120963, GSE121575, GSE122467, GSE123187, GSE128350, GSE129732, GSE130682, GSE135805, GSE137986, GSE142489, GSE143413, GSE144239, GSE147747, GSE149457, GSE151658, GSE151875, GSE151877, GSE151877, GSE152506, GSE153424, GSE153859, GSE154714, GSE156625, GSE156633, GSE156862, GSE158328, GSE158450, GSE158704, GSE158849, GSE159709, GSE160136, GSE161318, GSE161882, GSE161882, GSE163629, GSE164430, GSE164849, GSE165098, GSE166120, GSE166692, GSE166948, GSE167096, GSE167889, GSE169379, GSE169706, GSE169749, GSE171351, GSE171406, GSE171639, GSE173776, GSE174313, GSE178361, GSE178799, GSE178934, GSE179390, GSE179572, GSE180128, GSE181297, GSE182127, GSE182825, GSE182867, GSE182939, GSE184564, GSE185477, GSE186290, GSE188805, GSE188888, GSE189487, GSE189636, GSE190595, GSE190595, GSE192741, GSE193460, GSE194338; EMBL-EBI ArrayExpress (https://www.ebi.ac.uk/arrayexpress/) with accession number: E-MTAB-11114 and E-MTAB-9260; the 10x genomics website (https://support.10xgenomics.com/spatial-gene-expression/datasets/) and the SPATIAL research website (https://www.spatialresearch.org/resources-published-datasets/).

### Competing interests

We declare that one of our authors, Joshua Gould, is currently an employee of Cellarity.

### Funding

This work was supported by Genomics Data Center of Guangdong Province (No. 2021B1212100001) and Guangdong Provincial Key Laboratory of Genome Read and Write (No. 2017B030301011).

### Authors’ contributions

Conceptualization: XX, LQL, XFW, ZCX, and FZC; Data collection and curation: YH, ZCX, WWW and JC; Database construction: TY, TTL, WSD, FY, SJH, JC, and HL; Data analysis: ZCX, WWW, JC, and YH; Manuscript Draft: WWW. Manuscript review and editing: all authors; Dataset visualization: JG and JC; Project administration and supervision: XX, LQL, XFW, and FZC; Funding acquisition: XX, XFW, BW, and WJZ. All authors read and approved the final manuscript.

## Acknowledgements

We thank BGI Research for the technical support.

## Supplementary Figure

**Figure S1.**
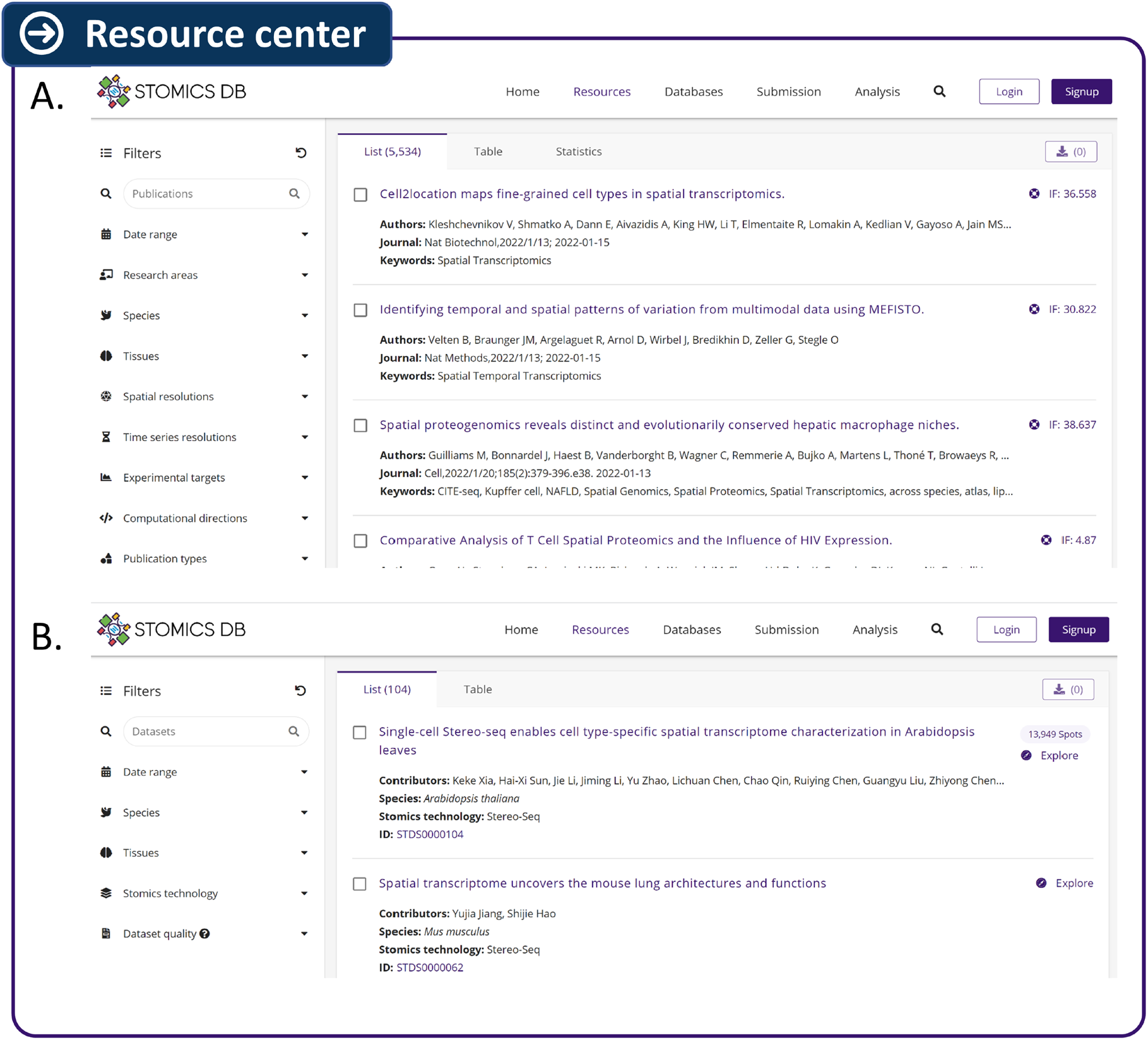
Interfaces of resource center of STOmicsDB. A. The interface of the publication search section. B. The interface of the dataset search section.

