## Supplementary figures and images for "STOmicsDB: a database of Spatial Transcriptomic data"

### Figure S1

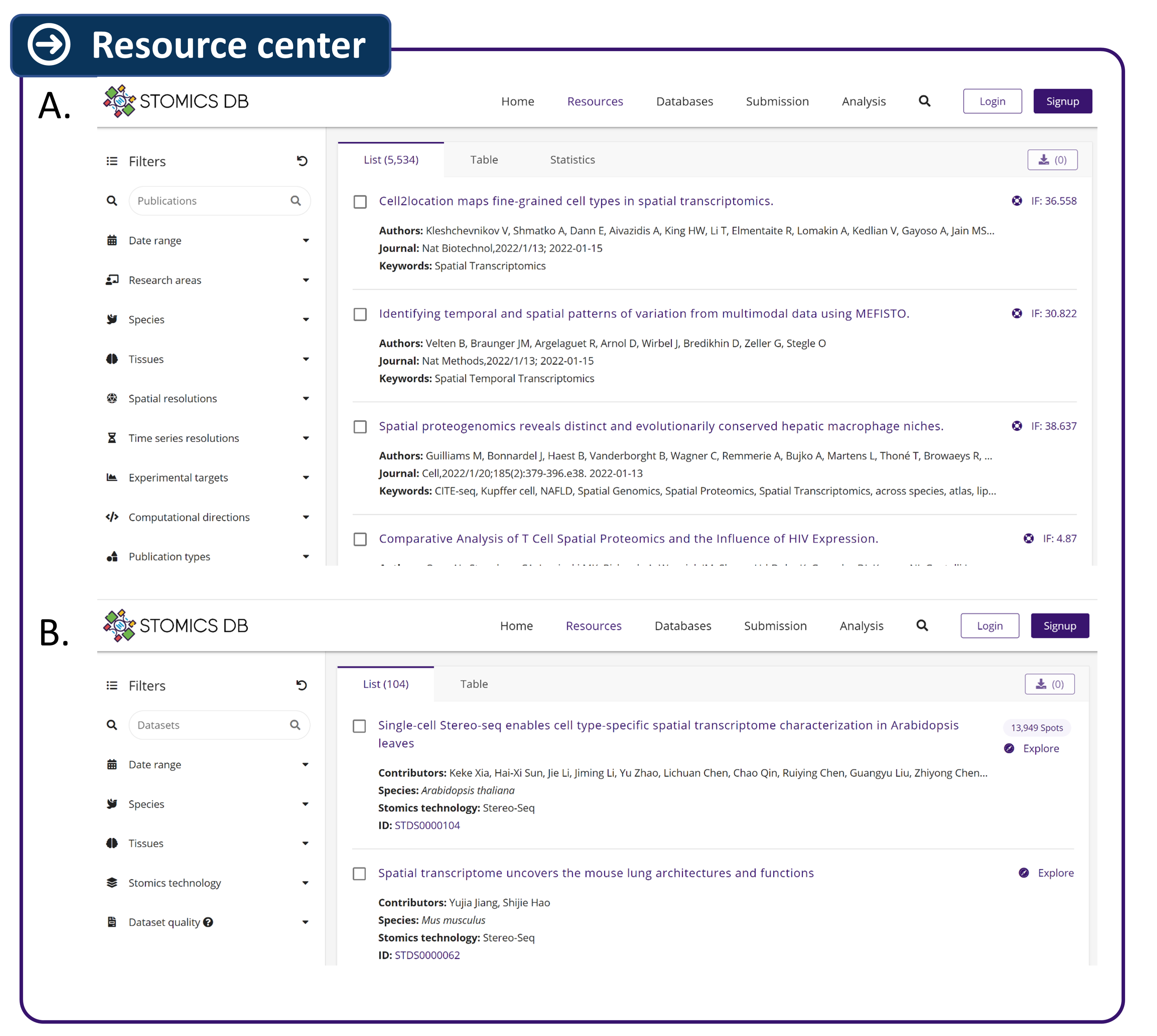
